# Insights into the bacterial profiles and resistome structures following severe 2018 flood in Kerala, South India

**DOI:** 10.1101/693820

**Authors:** Soumya Jaya Divakaran, Jamiema Sara Philip, Padma Chereddy, Sai Ravi Chandra Nori, Akshay Jaya Ganesh, Jiffy John, Shijulal Nelson-Sathi

**Author notes:** Author for Correspondence: Shijulal Nelson-Sathi, Computational Biology Laboratory, Interdisciplinary Biology, Rajiv Gandhi Centre for Biotechnology (RGCB), Thiruvananthapuram, India,.

## Abstract

Extreme flooding is one of the major risk factors for human health, and it can significantly influence the microbial communities and enhance the mobility of infectious disease agents within its affected areas. The flood crisis in 2018 was one of the severe natural calamities recorded in the southern state of India (Kerala) that significantly affected its economy and ecological habitat. We utilized a combination of shotgun metagenomics and bioinformatics approaches for understanding microbiome disruption and the dissemination of pathogenic and antibiotic-resistant bacteria on flooded sites. Here we report, altered bacterial profiles at the flooded sites having 77 significantly different bacterial genera in comparison with non-flooded mangrove settings. The flooded regions were heavily contaminated with faecal contamination indicators such as *Escherichia coli* and *Enterococcus faecalis* and resistant strains of *Pseudomonas aeruginosa, Salmonella* Typhi/Typhimurium*, Klebsiella pneumoniae, Vibrio cholerae* and *Staphylococcus aureus*. The resistome of the flooded sites contains 103 resistant genes, of which 38% are encoded in plasmids, where most of them are associated with pathogens. The presence of 6 pathogenic bacteria and its susceptibility to multiple antibiotics including ampicillin, chloramphenicol, kanamycin and tetracycline hydrochloride were confirmed in flooded and post-flooded sites using traditional culture-based analysis followed by 16S rRNA sequencing. Our results reveal altered bacterial profile following a devastating flood event with elevated levels of both faecal contamination indicators and resistant strains of pathogenic bacteria. The circulation of raw sewage from waste treatment settings and urban area might facilitate the spreading of pathogenic bacteria and resistant genes.

## Introduction

Flooding is one of the most destructive natural disasters, resulting in significant damages to life and infrastructure worldwide^1^. The devastating flooding in the southern state of India (Kerala) during August 2018 was declared as a “calamity of severe nature”, leaving 23 million people affected^2,3^. This was the worst ever flood in the history of Kerala since the Great flood in 1924^4^. During this season, Kerala state received a cumulative rainfall of 2346.3 mm, 42% greater than the monsoon average^5^. 35 out of 54 dams within the state were opened due to the heavy rainfall in its catchment areas. The flood left 10,319 houses fully damaged, more than 0.1 million houses partially damaged, destroyed 83,000 km of roads including 10,000 km of major roads, and 60,000 hectares of crops causing nearly $2.9 billion worth of damage^2^. Previous epidemiological evidence suggests that floods are positively associated with increased risk of water-borne and vector-borne diseases such as skin infection, typhoid fever, cholera, leptospirosis, hepatitis A, malaria, dengue fever, yellow fever and West Nile fever^6,7^. The stagnant floodwater can also significantly affect the environmental microbiome and spread of microbial pathogens^8^.

Previous studies showed floods due to Hurricane Katrina in 2005^9^, Chennai flood in 2015^10^, Hurricane Harvey in 2017^11^and Thailand flood in 2011^12^ significantly influenced the bacterial profile of water and soil. Reports on these floods evidenced that faecal contamination indicators like *Escherichia coli*, *Enterobacter aerogenes* and *Enterococcus* were widely distributed in water and soil sediments. In addition, infectious diseases causing pathogens such as *Legionella pneumophila*, *Vibrio cholerae, Aeromonas hydrophila*, *Klebsiella pneumonia, Clostridium perfringens, Salmonella Typhi*, *Streptococcus pyrogens* and *Shigella flexneri* were also abundant after the flood event. Another important concern is the possible coexistence of multidrug-resistant pathogenic bacteria with environmental bacteria, especially since the frequency of flooding could increase in coming years due to global climate change and expansion of coastal cities^13^. A previous study at Hurricane Harvey’s flooded sites in Houston revealed the elevated levels of anthropogenic antibiotic resistant markers such as *sul1* and *intI1*^11^. Furthermore, circulation of raw sewage from wastewater treatment plants and urban settings, have been shown to facilitate the spreading of antibiotic-resistant bacteria and resistant genes. There were attempts to understand flood associated microbial composition alteration^8,9,11^ and circulation of pathogenic and resistant bacteria, but its influence varies significantly based on the geography and nature of flooded environments. A better understanding of the impact of extreme flooding on the disruption of natural environmental microbiome and dissemination of pathogenic and antibiotic-resistant bacteria still needs detailed investigation. Kuttanad, an agricultural region in Kerala located in India’s lowest altitude of 4-10 feet below sea level is particularly susceptible to flood damage due to its unique geography. Here, we employed shotgun metagenomic and bioinformatics techniques to understand the bacterial community profile and the resistome of flood-affected areas of Kuttanad, India. We found a significant difference in the bacterial community structure and a higher abundance of multidrug-resistant pathogenic species in flood-affected areas. Our results will provide a better understanding of the health impacts of floods and provide more evidence to support decision-making for the prevention and control of flood-related diseases outbreaks.

## Results

Extreme flooding is one of the major risk factors for human health. It can significantly alter the top layer soil microbiome of flooded sites and enhance the mobility of infectious disease agents, especially the water-borne pathogens such as *Salmonella typhi*, *Vibrio cholerae*, *Leptospira sp.* and its resistant strains^15^. To investigate the bacterial community structure of flooded settings, we performed a detailed metagenomic screening of sediment samples collected from areas of Kuttanad (covers over 500 sq.km), devastated by heavy flooding (Fig. 1). Environmental indices were recorded at the sampling day (Supplementary Table S1).

**Figure 1:**
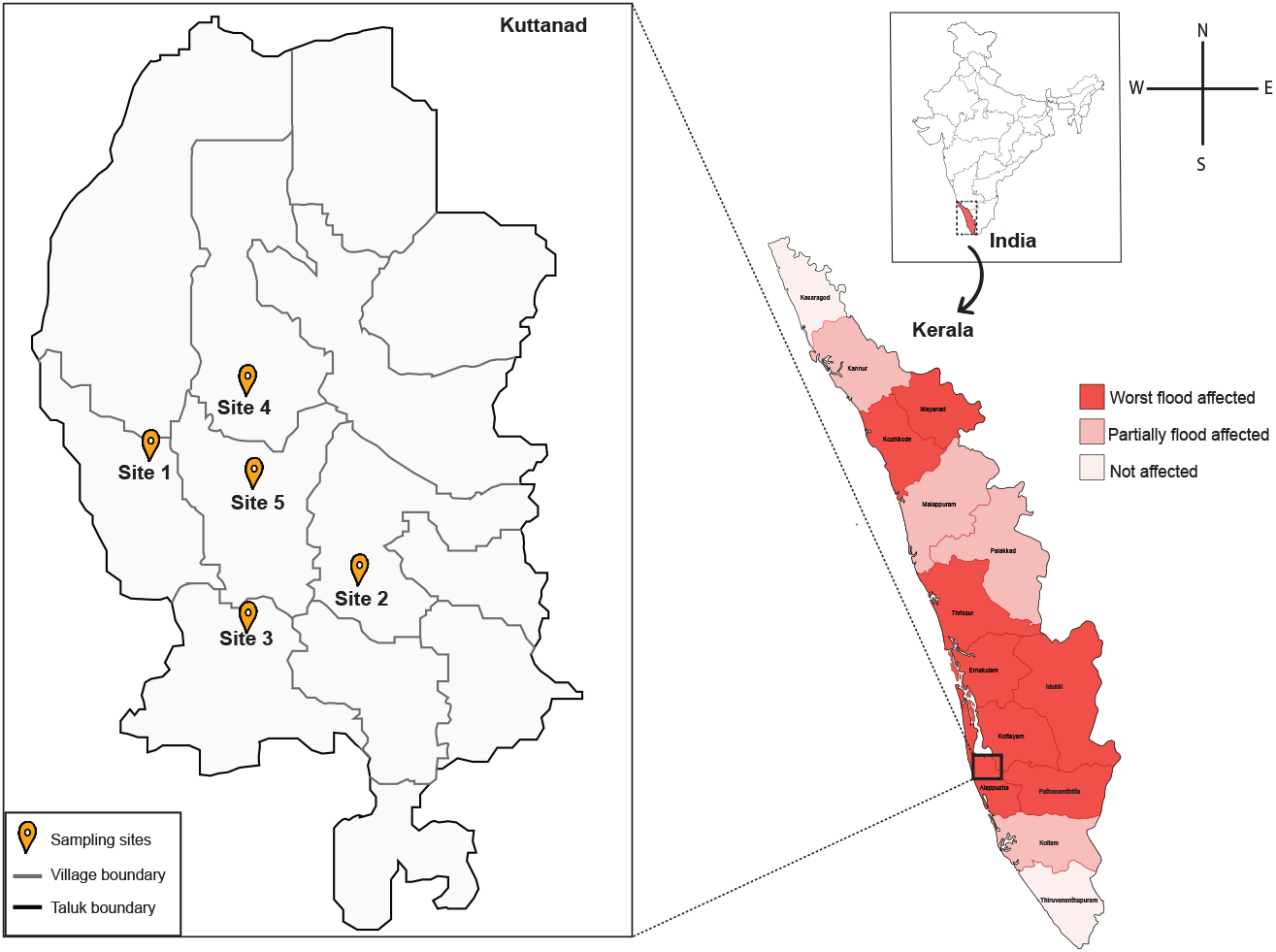
Map showing flooded regions and sampling sites of state Kerala, India. The intensity of the red colour indicates the level of flood severity in fourteen districts of the Kerala during August 2018. Triplicate samples were collected from each site during August 2018 and post-flood in February 2019.

### Bacterial profiles of flooded regions

On average, 178,527 16S rRNA reads were obtained from five different sites that were severely affected by the flood in August 2018. Less than half of the 16S rRNA reads (48 %) were could be taxonomically classified into known microbial organisms, of which the majority (96%) belongs to bacterial species. Among the annotated cases of bacterial species, Proteobacteria (45.4%) was found to be the most abundant phylum followed by Firmicutes (23%), Actinobacteria (16.39%) and Bacteriodetes (5.94%) (Fig. 2; Supplementary Table S2). Within the Proteobacterial phylum, the most abundant classes were the species of *Betaproteobacteria* (14.54%) and *Alphaproteobacteria* (14.42%) followed by *Gammaproteobacteria* (11%) and *Deltaproteobacteria* (4.73%). Within Firmicutes, the most abundant species were from *Bacilli* (12.96%) followed by *Clostridium* (9.18%) class. *Actinobacteria* (16.39%) was found to be the dominant class in the phylum Actinobacteria. The Bacteriodetes primarily consisted of *Bacteriodia* (4.15%) and *Flavobacteria* (1.27%). At the genus level, *Streptomyces* (6.03%) is the abundant genera followed by *Magnetospirillum* (4.08%), *Neisseria* (3.15%) and *Achromobacter* (2.66%).

**Figure 2:**
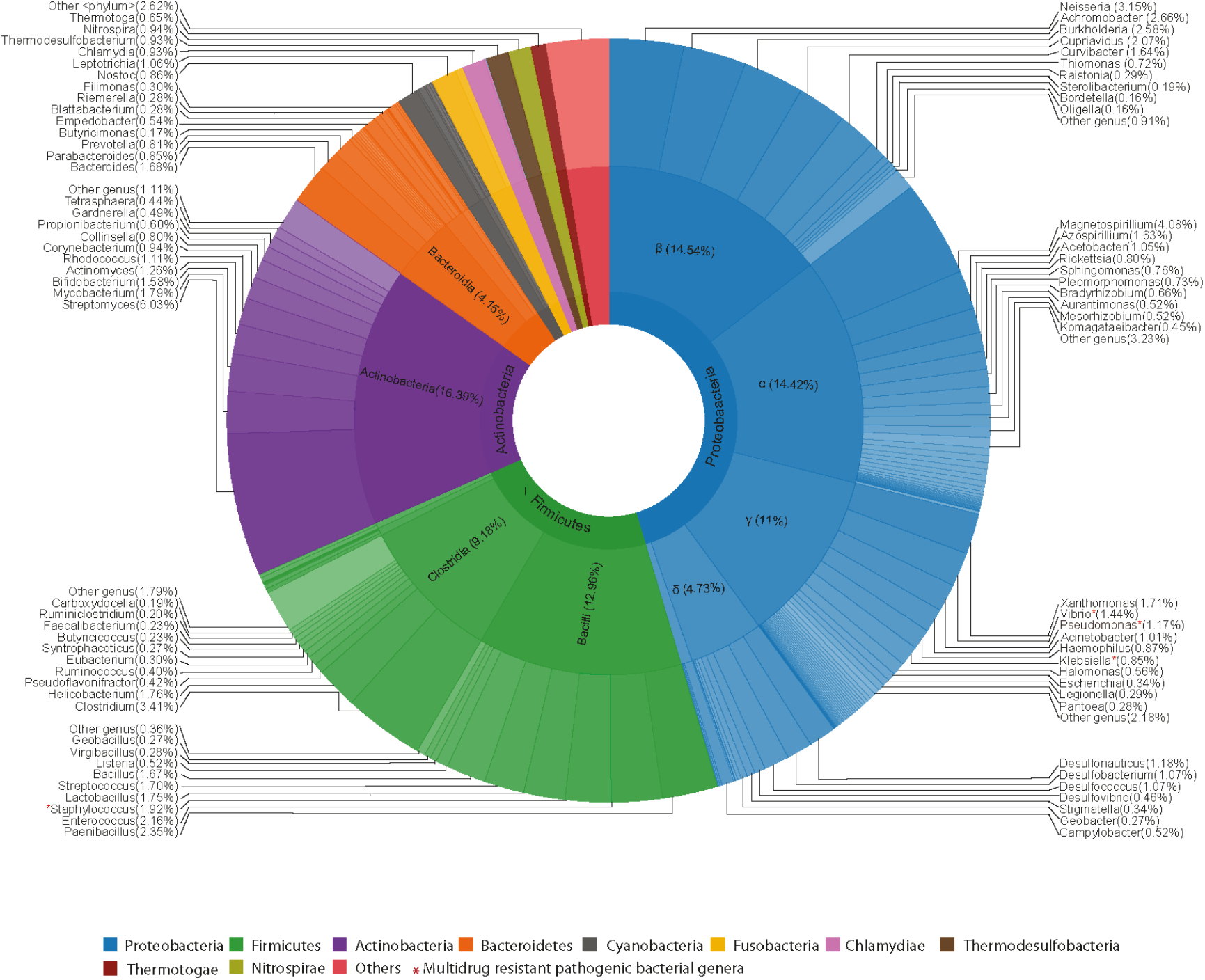
Bacterial taxonomic distribution of flooded sites. Sunburst plot showing the taxonomic classification and relative abundance of bacterial species in flood-affected regions. The Taxonomic phylum is represented in the innermost ring, class in the middle and genus represented in the outermost ring of the circle. Within each taxonomic classification, taxa are sorted according to its abundance. A * symbol represents the multidrug-resistant pathogenic bacteria present in flood-affected and post-flood affected regions. Taxa that were able to identify at least at the genus level are shown (Top ten genus). See Supplementary Table S2 for the full list of bacterial taxa found in flooded sites.

The bacterial community richness and diversity (chao1, Shannon indices) of flooded sites are found to be uniform, but its bacterial composition varies from non-flooded settings (Supplementary Table S3; Supplementary Fig. S1). A total of 77 genera were showing a significant difference (p<0.05) in their taxonomic distribution between flooded and non-flooded comparable mangrove settings. (Supplementary Fig. S2). Across multiple flooded sites, presence of *Streptomyces, Enterococcus, Magnetospirillum, Neisseria, Clostridium, Bacillus, Corynebacterium, Streptococcus, Helicobacterium* species were more abundant than non-flooded areas. The overall bacterial profile observed in the flooded sites in Kuttanad, Kerala was similar to the profile that is observed after Hurricane Harvey flood in Houston^11^.

Faecal contamination indicators and bacterial pathogens such as *Escherichia coli, Enterococcus faecalis, Klebsiella pneumoniae, Salmonella enterica* and *Legionella sp.* were detected in the flooded sites. The presence of *E. coli* and *Enterococcus* species in the flooded sites could increase the risk of eye, skin infections as well as oro-faecal infections such as diarrheal diseases, gastroenteritis ^16,17^. The increased faecal contamination levels clearly indicate that flood would elevate the spread of bacteria associated with anthropogenic sources.

### Resistome of flooded locations

To test the prevalence of Antibiotic Resistant Genes (ARGs) in the flooded sites, resistome profiles were reconstructed. Relatively similar resistant gene distribution was present, with an average number of 46 ARGs from all five flooded locations sampled. In total, resistome of flooded sites contains 103 unique genes that confer resistance to antibiotics over 12 different classes (Fig. 3). Number of ARGs present in flooded sites is 3 times higher compared to non-flooded sites for each class of antibiotics and only 27 ARGs (26%) were found to be common. Among the major resistant classes, most of the ARGs present in flood-affected sites confer resistance to aminoglycoside (19), beta-lactams (29), tetracycline (29), fluoroquinolone (31), macrolide (24) and phenicol (16). 38% of the detected ARGs in flooded sites were multidrug-resistant, the most frequent being *MexB*, *MexF* and *MuxB*. These genes are known to be encoded in plasmids and confer resistance against beta-lactams, fluoroquinolones, macrolides and phenicols.

**Figure 3:**
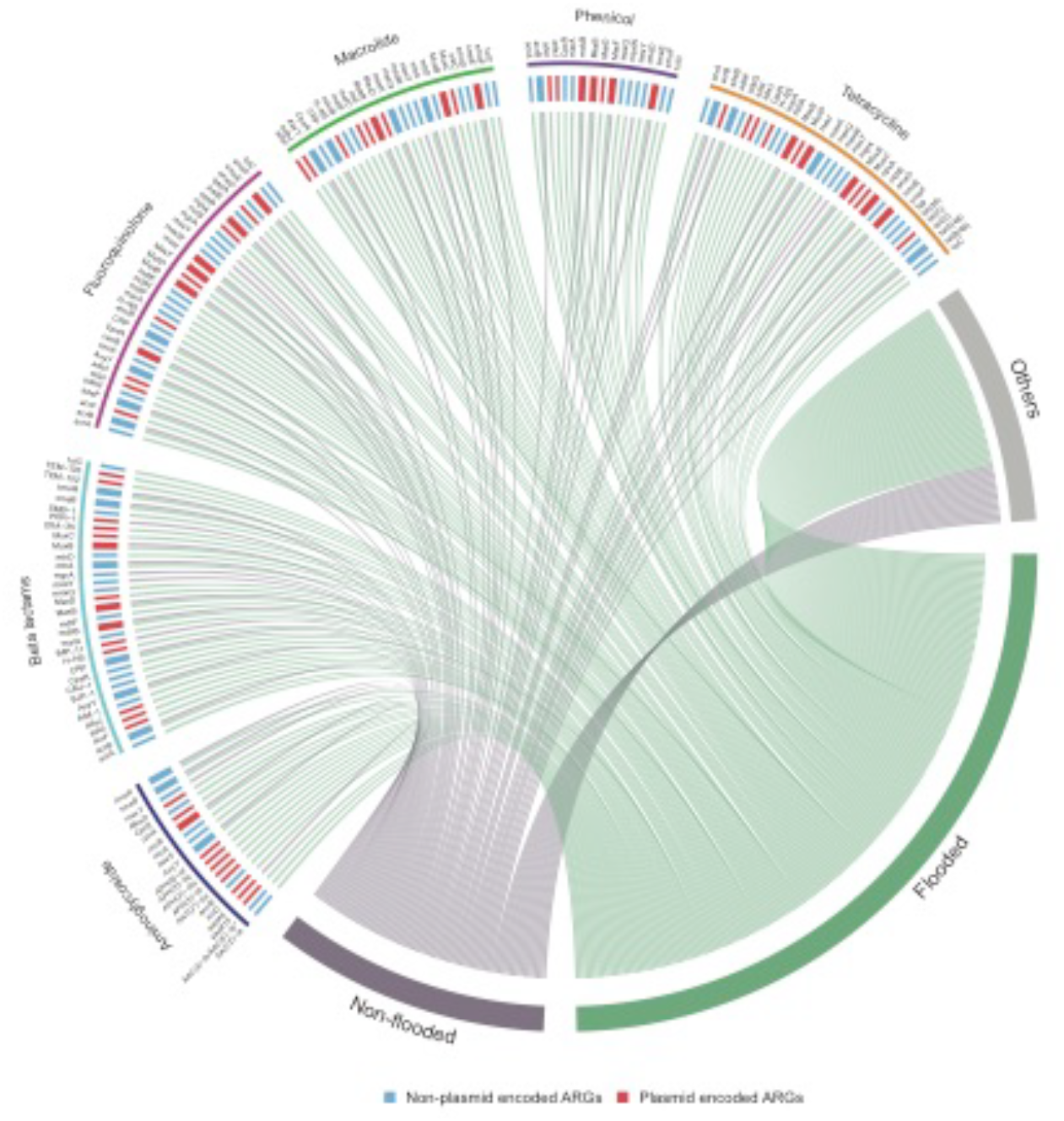
Resistome structure of flooded sites. Chord diagram showing the presence of Antibiotic Resistant Genes (ARG) detected in flooded (green) and non-flooded (grey) sites. Links are drawn when a specific ARG (small block) confers resistance to a certain drug class. The width of the gene block varies, according to it being shared or unique between samples. Red blocks indicate plasmid-encoded ARGs whereas blue blocks indicate non-plasmid encoded ARGs. ARGs conferring resistance to aminocoumarin, sulfonamide, mupirocin, rifampicin, triclosan, glycopeptide and diaminopyrimidine classes of antibiotics are represented as others category. Complete list of genes and its characteristics are listed in Supplementary Table S4.

Carbapenems, a subfamily of beta-lactam antibiotics currently is the most effective broad-spectrum antibiotics^18^, for which 10 resistant genes (eg., *mdsB*, *MexB, mexQ*) were found in the flooded sites. Additionally, we also detected genes that confer resistance to synthetic antibiotics such as sulfonamide (*sul1, sul2 and sul4*), fluoroquinolones (e.g., *smeE, adeF, acrB*) and penems (*TEM-126*and *TEM-102*). Furthermore, most of these genes were predominantly reported in species such as *Pseudomonas aeruginosa, Escherichia coli, Acinetobacter baumannii, Klebsiella pneumoniae, Shigella flexneri, Vibrio cholerae, Enterococcus faecalis, and Staphylococcus aureus* (Supplementary Table S4). In addition, we also found many virulence factors (VFs) in flooded sites that are distributed among the above bacterial pathogens. The functional classification of these VFs showed that they are involved in bacterial motility, cell adherence, iron uptake, secretions and toxins (Supplementary Table S5). These results indicate that flooded sites are large reservoirs of antibiotic-resistant genes and virulence factors associated with pathogenic mechanisms in clinically relevant bacteria.

The faecal bacterium population and ARG abundance might the result from the discharge of sewage from urban settings. There are about 39 (38%) resistant genes that are plasmid encoded, which increase the chances of conjugative transfer than non-ARG carrying plasmids. These genes are having a higher transfer potential, and while we could not precisely quantify the extent to which lateral gene transfer can promote further gene mobility, we can confidently report that the flood transported a significant number of resistant genes in the flooded sites.

### Pathogenic and resistant bacteria present in flooded settings are viable?

In order to check the viability and resistance of pathogenic bacteria present on flooded sites, we performed a culture-based analysis using selective and differential agar media, which includes Eosin methylene blue agar (EMB agar), Salmonella differential agar modified, HiCrome™Klebsiella selective agar base, Mannitol salt agar base, Thiosulphate-Citrate-Bile-Salt-sucrose agar (TCBS agar), Cetrimide agar base, Wilson-Blair agar with brilliant green (Wilson Blair agar w/BG) and Enterococcus differential agar base. Based on the morphology of bacterial colonies, we confirmed the presence of faecal contamination indicators such as *Enterococcus faecalis, Escherichia coli* and pathogenic bacterial species such as *Staphylococcus aureus, Salmonella* Typhi/Typhimurium*, Pseudomonas aeruginosa, Vibrio cholerae* and *Klebsiella pneumoniae*. The abundance of *Enterococcus faecalis* (8.4±0.5×10^3^ CFU/gram of dry weight) and *Escherichia coli* (3±0.35×10^3^ CFU/gram of dry weight) were 2 to 6 fold higher than comparable settings^19^ (Supplementary Table S6), but the abundance of *Enterococcus faecalis* was reduced to more than half (4100±0.4 CFU/gram of dry weight) in post-flood whereas *E.coli* was more or less similar in both during flood and post-flood. *Staphylococcus aureus*, an opportunistic pathogen was the most abundant bacteria with 8.5 ±3×10^5^ CFU/gram of dry weight in flooded samples and it was reduced to almost 2 fold in post-flood. Interestingly, the pathogenic species such as *Salmonella* Typhi/Typhimurium, *Vibrio cholerae*, *Klebsiella pneuomiae* and *Pseudomonas aeruginosa* are showing higher abundance in post-flooded samples compared to flooded samples. For example, *Salmonella* Typhi/Typhimurium and *Pseudomonas aeruginosa* were increased to 4.6 and 10.3 fold respectively.

The 6 pathogenic bacterial species identified in flooded sites were subjected to further antimicrobial susceptibility analysis against four different antibiotics such as ampicillin, chloramphenicol, kanamycin, and tetracycline hydrochloride. The bacterial isolates exhibited resistance against at least 3 antibiotics was considered as multidrug-resistant as proposed by Magiorakos *et al*., 2012^20^ and beta-lactams; aminoglycosides, phenicols and tetracyclines were on this category. The 20 resistant bacterial isolates showing characteristic colony morphology in selective media [such as yellowish green colonies (Cetrimide agar base), black with sheen colonies (Wilson Blair agar w/BG), purple-magenta coloured or cream to white coloured colonies (HiCrome™Klebsiella selective agar base), pink to red colonies (Salmonella differential agar modified), round yellow colonies (TCBS agar), yellow or white colonies surrounded by a yellow zone (Mannitol salt agar) and purple with black center and green metallic sheen colonies(EMB agar)] were randomly picked up and their 16S rRNA genes were sequenced using universal primers. Interestingly, we found that these 20 isolates belong to species such as *Pseudomonas aeruginosa*, *Salmonella* Typhi/Typhimurium*, Klebsiella pneumoniae, Vibrio cholerae, Staphylococcus aureus* and *Escherichia coli.* Among these, 2/2 isolates of *Pseudomonas aeruginosa* and 2/3 isolates of *Salmonella* Typhi/Typhimurium were resistant to all four antibiotics tested in flood and post-flood samples. In addition, 2/7 isolates of *Klebsiella pneumoniae* and 2/4 isolates of *Vibrio cholerae* were multidrug-resistant in flood and post-flood samples (Fig. 4 and Fig. 5). *Staphylococcus aureus* was sensitive to all four antibiotics during the flood and resistant in post-flood while *Escherichia coli* were resistant to only ampicillin in flood and post-flood samples.

**Figure 4:**
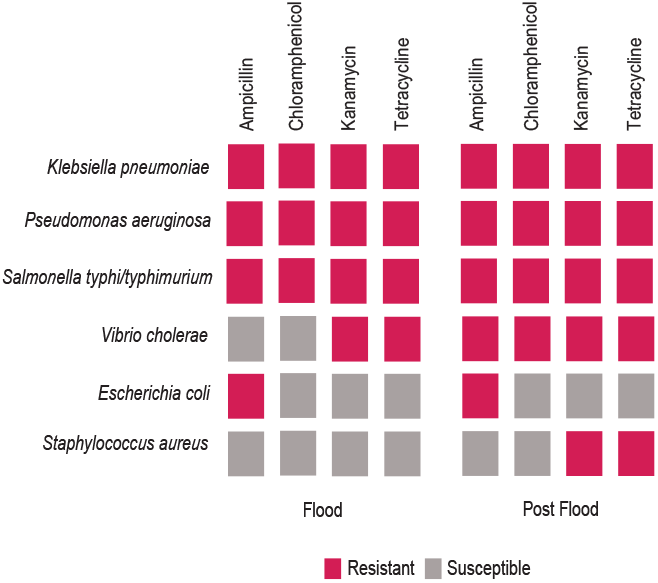
Antibiotic susceptibility profile of pathogenic bacterial species isolated from flooded and post-flood sites. Red colour indicates resistance against antibiotics whereas grey colour indicates susceptible to the antibiotic.

**Figure 5:**
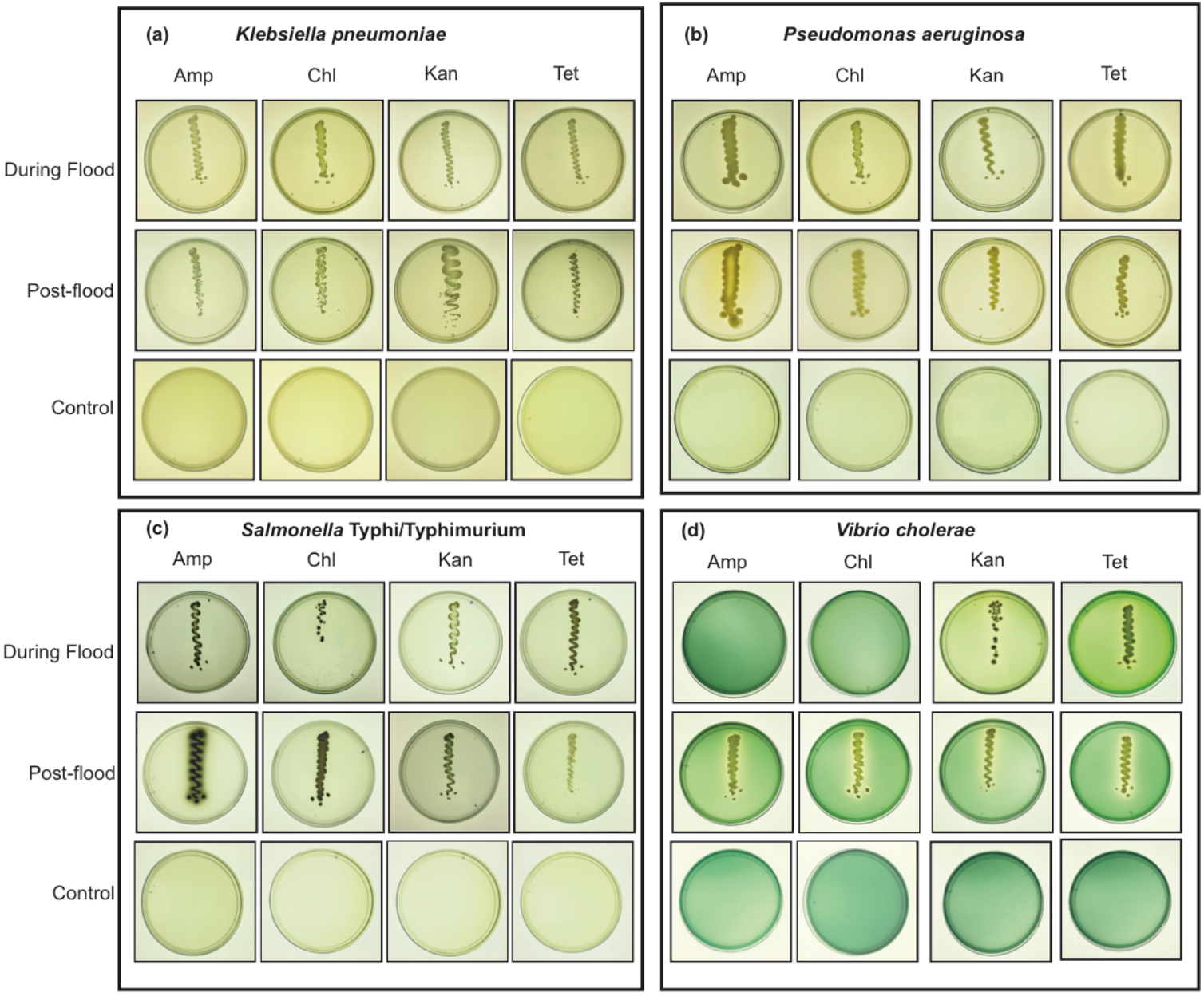
In vitro evaluation of antimicrobial resistance of pathogenic bacterial species isolated from flooded sites (during flood and post-flood). The images showing pathogenic bacteria streaked on selective/differential agar media containing different antibiotics such as Amp: Ampicillin (100 µg/ml), Kan: Kanamycin (50 µg/ml), Chl: Chloramphenicol (25 µg/ml), Tet: Tetracycline hydrochloride (10 µg/ml) and incubated at 37ºC for 1-3 days. (a) *Klebsiella pneumoniae* (HiCrome™ Klebsiella selective agar base). (b) *Pseudomonas aeruginosa* (Cetrimide agar base). (c) *Salmonella* Typhi/Typhimurium (Wilson Blair agar w/BG). (d) *Vibrio cholerae* (TCBS agar).

## Discussion

Our results demonstrate that the devastating flood that occurred in the southern state of India significantly influenced the bacterial composition of its watershed areas. Shotgun metagenomic profiling revealed significantly altered bacterial profile at the flooded sites. However, a large portion (52%) of the microbial diversity of these flooded settings still remain unknown. Most of the microbes detected are environmental bacteria but faecal contamination indicators and clinically relevant pathogens such as *Vibrio cholerae, Klebsiella pneumoniae, Salmonella* Typhi/Typhimurium etc and its resistant strains were also abundant. The higher levels of bacterial contamination and dissemination of resistant pathogenic bacteria at the flooded areas might cause water-borne and vector-borne diseases^21^ such as dysentery, cholera, typhoid fevers and other gastrointestinal diseases.

As previously reported by Garner E *et al*., a large number of resistant genes, which shows resistance to major classes of antibiotics were observed in flooded sites in Colorado, 2013^22^. Our study also finds a similar number of multidrug-resistant genes at the flooded sites and the most common mechanism of resistance as efflux pump. Further, we found a high prevalence of resistant genes that are associated with clinically relevant pathogens and their viability was confirmed by culture-based techniques. The coexistence of bacterial species of different resistant level could increase the chance of resistant genes being exchanged between strains of pathogenic and non-pathogenic bacteria^23^. 46% of the multidrug-resistant genes identified were found to be plasmid encoded, which increase the transfer potential of these genes ^24,23^. Furthermore, better time-resolved sampling is required to estimate pathogen survival duration and source of detected pathogens and resistant strains and their evolution ^25^.

The higher abundance of bacterial pathogens carrying multidrug-resistant genes is alarming because it could make post-flood disease outbreaks difficult to treat. Our study provides more insights into the pathogenic and resistance traits of bacterial communities of flooded and post-flooded settings, which helps to plan better preventive measures against post-flood disease outbreaks; preventive measures such as chlorination of floodwater, vaccination and good hygienic practices that can help avoid the spread of infectious diseases.

## Materials and methods

### Sample collection

Soil/sediment samples were collected from five sites of flood-affected areas in Kuttanad, Kerala namely Nedumudi (9°26’33.73” N, 76°24’26.565” E), Ramankary (9°24’47.808’’ N, 76°27’19.152” E), Thakazhy (9°23’5.352 N, 76°26’42.971” E), Pulinkunnu (9°26’55.464 N”, 76°26’47.76” E), Mankombu (9°25’19.056” N, 76°28’19.92” E). The sampling was performed during the flood season and post 6 months of flooding (August 2018 and February 2019). As sampling sites were public places, no need for particular permission was required for samples collection. Triplicate sediment samples were collected from each place in sterile 50ml conical tubes by using sterile steel scoops, and unique identifiers were given for each sample. Geographical location and environmental indices were recorded at the time of sampling. PH meter in a suspension of 1:5 ratios of soil and ultrapure water measured the pH of the soil sediment samples at the day of sampling. Following sample collection, 50ml conical tubes were wrapped with parafilm and transported to the laboratory on ice. The sediment samples were stored in the deep freezer for DNA extraction and microbial analysis.

### DNA Extraction

Metagenomic DNA from sediment samples was extracted using Power soil DNA extraction kit (MO Bio Laboratories) and slight modifications were made in the Metagenomic procedure such as the addition of RNase A (1µg/ml, Qiagen) along with solution C2 (inhibitor removal solution), 1h incubation at 37°C, and two additional washes were performed with 70% ethanol^26^. The purity and concentration of the metagenomic DNA was measured using multimode microplate reader (TecanSpark 10m, Tecan, Switzerland). The integrity of extracted DNA was confirmed by agarose gel electrophoresis by loading equal amount of extracted DNA on agarose gel along with גHindIII digest marker (NEB) ^27^. The DNA was pooled from subsamples from each site and used for shotgun metagenomic sequencing.

### Shotgun metagenomic sequencing and analysis

One microgram high quality metagenomic DNA was further used for shotgun metagenomic sequencing. Metagenome sequencing libraries were prepared using the NEBNext Ultra DNA Library preparation kit following the manufacturer’s protocol. In brief, the DNA is subjected to a sequence of enzymatic steps for repairing the ends and tailing with dA-tail followed by ligation of adapter sequences. These adapter-ligated fragments were then cleaned up using SPRI beads. The cleaned fragments were indexed using limited PCR cycle to generate final libraries for paired-end sequencing. 2×150bp sequencing reads were generated on the Illumina HiSeq system, yielding about 3 GB of data per sample.

For comparative analysis, we used 4 metagenomic datasets (SRR2844600, SRR2844601, SRR2844602, SRR2844616) of sediment samples collected from mangrove settings reported by Imchen M *et al.* in 2017^28^. The quality of paired-end sequences was assessed using FastQC v0.11.4^29^. Reads with low quality (Qvalue20 cutoff) and adapter sequences were trimmed and removed using FastX toolkit v.0.0.13.2 ^30^. SortMeRNA^31^ was applied to the filtered shotgun metagenome data to extract 16S rRNA sequences. For each 16S rRNA sequence, a BLASTx search was performed against the NCBI non-redundant protein database using DIAMOND v.0.9.24^32^. Output data were analyzed with the MEGAN5 analyzer v5.7.1^33^, using the following settings: Min Score = 50, Top Percent = 10, Min Support = 1, Min-Complexity Filter = 0. Comparative analysis for taxa in terms of percentage mean relative frequency was performed using STAMP V2.1.3^34^, G-test (w/Yates’) + Fisher’s was applied to compare bacterial communities pairwise (Mangrove and Kuttanad) with *P* value <0.05, confidence intervals of 95% and extended error bars were plotted. Genus-level taxa abundances were used in STAMP^34^ to generate principal component analyses (PCA). Alpha diversities were measured by Shannon diversity index and chao1 index using Qiime2^35^to analyze the diversity within the samples.

Antibiotic resistance and virulence genes were identified by mapping the quality filtered reads against CARD^36^ and VFDB ^37^using DIAMOND v.0.9.24 (BLASTx, -e 1e-05). Only hits with sequence identity above 90% and an alignment length over 25 amino acids were kept ^38,39^. The identified ARGs were further annotated according to the corresponding CARD database descriptions.

The occurrence of ARGs in plasmids was determined using BLASTn^40^ against 14,595 complete plasmid sequences from the NCBI RefSeq database (updated on 16 May 2019). Hits with >70% of query coverage and >70% of amino acidic identity was kept.

### Abundance of selected pathogens

The abundance of selected pathogenic bacteria was calculated by colony forming unit. One gram of soil sediment sample was pooled from flood-affected areas and mixed with 9mL sterile phosphate buffered saline taken in test tubes. The solution was homogenized by using a vortex machine and the diluted suspension was serially diluted up to 10^−4^ ^41^. 100µL of diluted suspension from each dilution was spread on different selective media such as Eosin Methylene Blue agar (EMB Agar, HiMedia), Salmonella differential agar modified (HiMedia), HiCrome™Klebsiella selective agar base (HiMedia), Mannitol salt agar (HiMedia), Thiosulphate-Citrate-Bile-Salt-sucrose agar (TCBS agar, HiMedia), Cetrimide agar base (HiMedia), Wilson-Blair agar with brilliant green (WB agar w/BG, HiMedia), Enterococcus differential agar base (HiMedia), by using a sterilized glass spreader and incubated at 37°C for 1-3 days depending upon the bacteria that were grown in the selective media^42^. Control plates were spread with only phosphate buffered saline and processed in parallel with soil suspension as method blank. To minimize error, all these experiments were repeated thrice. After the incubation period, the numbers of colonies from each plate were counted and CFU of wet soil was calculated by the following equation.

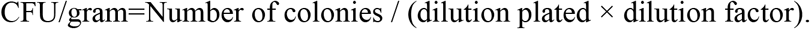

Colony forming unit of soil bacteria are expressed in dry weight instead of one gram of wet soil. So, one gram of sediment samples were pooled from flood-affected areas and then air-dried to determine the percentage change. Percentage change is calculated by the following equation and this percentage change value is used to convert CFU per gram-wet soil to CFU per gram dry soil. Experiment is performed thrice for the sediment samples collected during flood and post-flood, result is expressed in CFU (mean ± s.d)

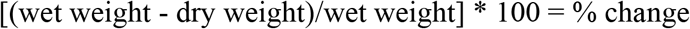

### Identification of antibiotic resistant bacterial pathogens

Bacterial pathogens that were grown on selective media and were used to check resistance against ampicillin (Sigma-Aldrich), chloramphenicol (Sigma-Aldrich), kanamycin (Sigma-Aldrich) and tetracycline (Sigma-Aldrich). Pure colonies were picked from the selective media and incubate in 5ml nutrient broth for 1-3 days at 37°C for 200rpm in Innova shaker (New Brunswick™Innova (®40/40R). After incubation, a loop full of bacterial culture was streaked on respective selective media containing antibiotics such as ampicillin (100µg/mL), chloramphenicol (25µg/mL), kanamycin (50µg/mL), and tetracycline hydrochloride (10µg/mL) respectively. The plates were incubated at 37°C for 1-3 days according to the bacteria that streaked on the media. A loop full of nutrient broth streaked on the antibiotic containing media is used as a method blank and processed parallel with bacterial culture.

### 16S rRNA Sequencing

Bacteria showing antibiotic resistance were picked from the selective media and incubated in 10ml LB broth for 12 hrs. After incubation, bacterial genomic DNA was isolated using the Nucleospin®Microbial DNA Kit (Macherey Nagel) according to the manufacturer’s protocol. The quality and concentration of genomic DNA was measured by using a multimode microplate reader (Tecan Spark 10m, Tecan, Switzerland) and high-quality genomic DNA was used for 16S sequencing. 16S rRNA gene was amplified using a universal forward primer (5’CAGGCCTAACACATGCAAGTC3’) and reverse primer (3’GGGCGGWGTGTACAAGGC5’) ^43^. PCR was carried out in a 20 µl reaction volume which contained 1X PCR buffer (100mM Tris HCl, pH-8.3; 500mM KCl), 0.2mM each dNTPs, 2.5mM MgCl_2_, 1 unit of AmpliTaq Gold DNA polymerase enzyme (Applied Biosystems), 0.1 mg/ml BSA, 4% DMSO, 5 pM of forward and reverse primers and template DNA in GeneAmp PCR System 9700, Applied Biosystems). PCR conditions were; 1 cycle of 95 °C for 5 minutes followed by 35 cycles of 95 °C for 30s, 65 °C for 40s, 72 °C for 60s and 72°C for 7minute. The PCR product was checked in agarose gel electrophoresis by loading 5µl of PCR product on to 1.2% (w/v) agarose gel and the gel was visualized using UV transilluminator (Genei) and the image was captured using Gel documentation system (Bio-Rad). Amplified products were re-purified by using ExoSap (Thermo Fischer Scientific) treatment. Sequencing PCR was carried out by using the BigDye Terminator v3.1 Cycle sequencing Kit (Applied Biosystems, USA) in a thermal cycler (GeneAmp PCR System 9700, Applied Biosystems) according to the manufacturer’s instruction^44^.The forward and reverse sequences obtained from Sanger sequencing were merged using Bioedit v7.1^45^. Further, the merged sequences were blasted against the reference data sets of NCBI for identification of species.

### Data availability

The raw metagenomic data reported in this paper have been deposited in the NCBI Sequence Read Archive with accession numbers SRR9620086, SRR9620087, SRR9620088, SRR9620089, SRR9620090, as part of BioProject PRJNA552210 (NCBI Bio-Project database *http://www.ncbi.nlm.nih.gov/bioproject*).

## Acknowledgements

We thank Prof. M. Radhakrishna Pillai, and Dr. TR Santhosh Kumar, Rajiv Gandhi Centre for Biotechnology (RGCB), Kerala, India for providing infrastructure support for conducting this study. This work was supported by YIPB program of Kerala State Council for Science, Technology and Environment (KSCSTE) [043/YIPB/KBC/2017/KSCSTE]; and INSPIRE Faculty Fellowship to S.N.-S [DST/INSPIRE/04/2015/002935]; and Early Career Award [ECR/2017/002980] by the Department of Science and Technology.

## Author Contributions

S.N.-S designed the study. S.J.D, J.S.P, J.J, S.R.N collected all sediment samples. S.J.D and P.C performed the culture-based analysis. J.S.P performed the metagenomic data analysis with assistance from S.R.N and A.J.G. S.N.-S wrote the manuscript with input from all authors. All authors discussed the results and contributed to the final manuscript.

