## Supplementary Figures and Tables for "Insights into the bacterial profiles and resistome structures following severe 2018 flood in Kerala, South India": Supplementary Table S1.docx

Supplementary Table S1: Table showing the geographical details of sampling sites and its environmental indices during the flood (August 2018) and post-flood (February 2019) in Kuttanad, Kerala, India.

| **Sampling site** | **Sample code** | **Name of site** | **Coordinates** | **Date of sample collection** | **Temperature** | **Weather**  **condition** | | **Humidity** | **pH** |
| --- | --- | --- | --- | --- | --- | --- | --- | --- | --- |
| **During flood (August 2018)** | | | | | | | | | |
| Site 1 | RGCB 1027 | Nedumudi | N9°26’33.73”  E76°24’26.565” | 29-08-2018 | 25°C | Cloudy | | 90% | 6 |
| Site 2 | RGCB 1028 | Ramankary | N9°24’47.808”  E76°27’19.152” | 29-08-2018 | 25°C | Cloudy | | 90% | 6 |
| Site 3 | RGCB 1029 | Thakazhy | N9°23’5.352 E76°26’42.971” | 29-08-2018 | 26°C | Cloudy | | 86% | 5 |
| Site 4 | RGCB 1030 | Pulinkunnu | N9°26’55.464” E76°26’47.76” | 29-08-2018 | 27°C | Cloudy | | 85% | 6 |
| Site 5 | RGCB 1031 | Mankombu | N9°25’19.056” E76°28’19.92” | 29-08-2018 | 27°C | Cloudy | | 84% | 6 |
| **Post-flood (February 2019)** | | | | | | | | | |
| Site 1 | RGCB 2027 | Nedumudi | N9°26’33.73”  E76°24’26.565” | 07-02-019 | 26°C | Partially cloudy | 80% | | 5.5 |
| Site 2 | RGCB 2028 | Ramankary | N9°24’47.808”  E76°27’19.152” | 07-02-019 | 29°C | Clear | 65% | | 6 |
| Site 3 | RGCB 2029 | Thakazhy | N9°23’5.352 E76°26’42.971” | 07-02-019 | 30°C | Partially cloudy | 61% | | 5 |
| Site 4 | RGCB 2030 | Pulinkunnu | N9°26’55.464” E76°26’47.76” | 07-02-019 | 28°C | Humid & Partially cloudy | 72% | | 5.8 |
| Site 5 | RGCB 2031 | Mankombu | N9°25’19.056” E76°28’19.92” | 07-02-019 | 31°C | Partially cloudy | 57% | | 5 |
