## Supplementary Figures and Tables for "Insights into the bacterial profiles and resistome structures following severe 2018 flood in Kerala, South India": Supplementary Table S3.docx

| Sample ID | Shannon diversity index, H’ | Chao1 richness |
| --- | --- | --- |
| Flooded samples | | |
| RGCB_1027 | 6.13 | 188 |
| RGCB_1028 | 6.23 | 278 |
| RGCB_1029 | 6.19 | 169 |
| RGCB_1030 | 6.54 | 271 |
| RGCB_1031 | 6.27 | 172 |
| Non-flooded samples | | |
| SRR2844600 | 6.23 | 304 |
| SRR2844601 | 6.29 | 174 |
| SRR2844602 | 6.52 | 284 |
| SRR2844616 | 5.03 | 164 |

Supplementary Table 2: Table showing the biodiversity index of bacterial communities in flooded and non-flooded sites.
