## Supplementary Figures and Tables for "Insights into the bacterial profiles and resistome structures following severe 2018 flood in Kerala, South India": Supplementary Table S5.docx

Supplementary Table 5: Functional annotation of the virulence factors distributed in pathogenic species found in flooded sites

| **Functions** | Pathogenic Species | | | | | | |
| --- | --- | --- | --- | --- | --- | --- | --- |
|  | ***Acinetobacter baumannii*** | ***Escherichia coli*** | ***Klebsiella pneumoniae*** | ***Pseudomonas aeruginosa*** | ***Salmonella enterica*** | ***Staphylococcus aureus*** | ***Vibrio cholerae*** |
| **Adherence** | ompA | yagZ/ecpA, yagX/ecpC | fimD, fimA | fleQ, flgG, flhA, flgI, fliC, fliF, fliG, fliI, fliM, fliN, fliP, fliI, waaG, chpA, chpE, pilA, pilB, pilC, xcpA/pilD, pilR, pilT, pilU, pilG, pilJ | csgG, fimD, flhC, fliI, fliM |  | tcpI, pilB |
| **Secretion**  **System** |  | espL1 | clpV/tssH, hcp/tssD, icmF/tssM, tssF,  vgrG/tssI, | xcpS, xcpR |  |  | clpB/vasG,  vipB/  mglB, epsE |
| **Toxin** |  | hlyE/clyA, hlyB |  |  |  |  |  |
| **Biofilm**  **Formation** | adeF, adeG, pgaC |  | mrkD, mrkB |  |  |  |  |
| **Enzyme** | plc |  | acrA, acrB |  |  |  |  |
| **Immune**  **Evasion** | pgi |  | gmd, gnd, manB, ugd  WcaG, |  |  |  |  |
| **Iron uptake** | basB, basD | chuA | iutA, entC, entA, entE, entF,  A225_1607, fepD, fepB, | pvdD, pvdI |  |  |  |
| **Regulation** | bfmR |  | rcsB |  | fur, phoP, rpoS |  |  |
| **Serum resistance** | ompA, pbpG |  |  |  |  |  |  |
| **Motility** |  |  |  |  | cheY, cheB, cheR, tar/cheM, |  | flrA, cheY |
| **Antiphagocytosis** |  |  |  | algB, algR,  algI, algA, mucD |  | cap8F, cap8G | cpsA, wbfY |
| **Magnesium**  **Uptake** |  |  |  |  | mgtB |  |  |
| **Nutritional factor** |  |  | allC |  |  |  |  |
| **Total VF** | 45 | 43 | 48 | 69 | 18 | 2 | 9 |
