## Supplementary Figures and Tables for "Insights into the bacterial profiles and resistome structures following severe 2018 flood in Kerala, South India": Supplementary Table S6.docx

| Microorganism | During flood  (CFU / gram of dry weight) | Post-flood  (CFU/gram of dry weight) |
| --- | --- | --- |
| *Staphylococcus aureus* (10^5^) | 8.5±3.0 | 4.3±0.1 |
| *Enterococcus faecalis* (10^3^) | 8.4±0.5 | 4.1±0.4 |
| *Escherichia coli* (10^3^) | 3±0.35 | 3±0.15 |
| *Salmonella* Typhi/Typhimurium(10^3^) | 12.03±4 | 56.3±2.5 |
| *Vibrio cholera* (10^3^) | 2.5±1 | 3.5±1.36 |
| *Klebsiella pneumoniae* (10^3^) | 1.8±1.36 | 5.8±0.64 |
| *Pseudomonas aeruginosa* (10^2^) | 3.3±1.44 | 34±1.65 |

Supplementary Table S6: Table showing the Colony forming unit (CFU) of faecal indicator bacteria and pathogenic bacteria in soil/sediment samples collected during the flood (August 2018) and post-flood (February 2019). Abundance of *Escherichia coli*, *Enterococcus faecalis*, *Vibrio chloerae*, *Klebsiella pneumoniae*, *Staphylococcus aureus*, *Pseudomonas aeruginosa* and Salmonella Typhi/Typhimurium are represented in CFU/gram of dry weight*.*

All data are mean of triplicates; ±standard deviation (s.d).
