## Supplementary figures and images for "Insights into the bacterial profiles and resistome structures following severe 2018 flood in Kerala, South India"

### Supplementary Figure S1.pdf

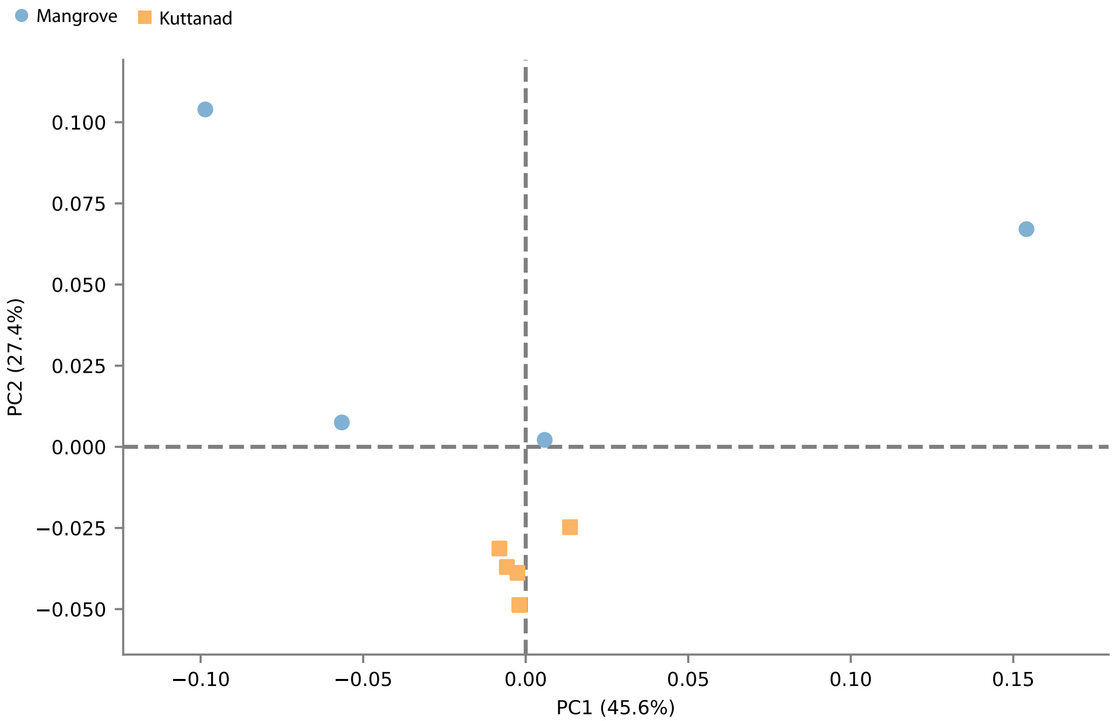

### Supplementary Figure S2.pdf

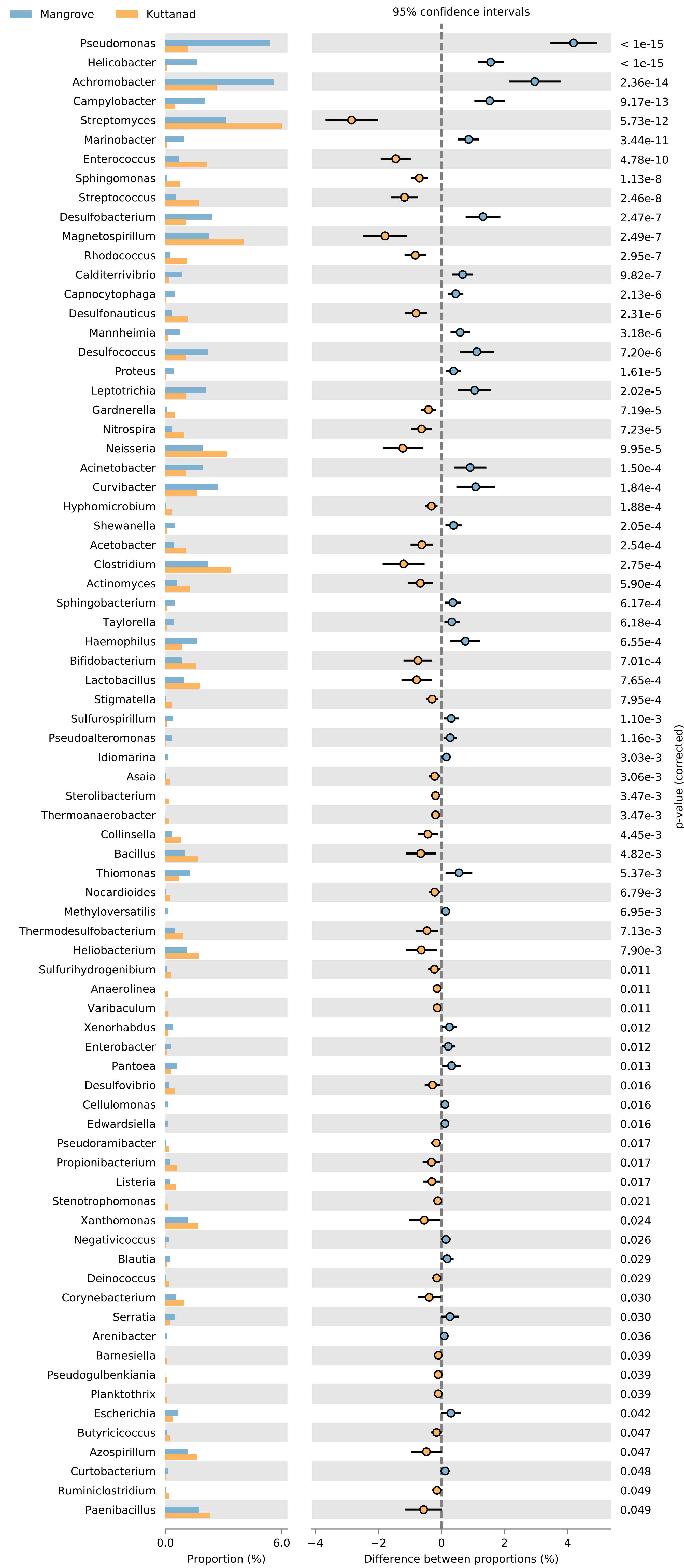
